# Plastidial Phosphorylase (Pho1a) is the dominant glucosyltransferase regulating starch granule initiation in potato tubers

**DOI:** 10.64898/2026.08.24.746836

**Authors:** Ciara O’Brien, Matthew Carswell, Aaron Rowland, Danielle Scarbrough, Xintong Huang, Brendan Fahy, Joerg Fettke, Jeffrey W. Habig, David Seung

## Abstract

Starch granule initiation involves the extension of maltooligosaccharide primers by glucosyltransferases. STARCH SYNTHASE 4 (SS4) plays a central role in almost all examined plant species, while the plastidial PHOSPHORYLASE 1 (Pho1) also plays an important role in some species, including rice and wheat. In Arabidopsis, an additional enzymatically inactive homolog of SS4, STARCH SYNTHASE 5 (SS5) contributes to starch granule initiation. To elucidate the mechanism of starch granule initiation in potato tubers, we used CRISPR/Cas9 to generate *ss4*, *ss5*, and *pho1a* knockout mutants in the commercial tetraploid ‘Clearwater Russet’, to systematically investigate their contribution to granule initiation. In *ss4* and *ss5* tubers, starch granule size and morphology were unaltered relative to the wild type, suggesting that SS4 and SS5 are dispensable for normal granule initiation in potato tubers. In contrast, *pho1a* tubers had compound starch granules that arose from multiple initiations, greatly reduced granule size, and highly variable granule morphologies. Affinity pull-down to find Pho1a interaction partners identified LIKE EARLY STARVATION (LESV), although yeast 2-hybrid assays did not show direct protein-protein binding. When expressed alone in *Nicotiana benthamiana* leaves, Pho1a located to the chloroplast stroma, but when expressed alongside LESV, both proteins co-located on starch granules. This co-localisation, alongside the similar accumulation of small starch granules when LESV is knocked out in tubers, suggest a possible functional interaction *in planta*. These findings position Pho1a as the central glucosyltransferase in starch granule initiation in Clearwater Russet tubers, where it acts together with LESV.

**Significance statement:** Through knocking out candidate glycosyltransferases implicated in starch granule initiation in potato, we found *pho1a* mutants had altered number, size, and shape of granules, positioning Pho1a as a central regulator of granule initiation in potato. Surprisingly, *ss4* mutants showed no change in granule initiation, contrasting results in Arabidopsis and cereals, and demonstrating diverse mechanisms of starch initiation among staple crops.

## Introduction

Starch is a major form of carbohydrate storage in plants, which is degraded into soluble sugars to fuel growth and metabolism in the absence of photosynthesis. Transitory starch accumulates in chloroplasts and is degraded at night to sustain autotrophic growth. Heterotrophic organs, including potato tuber, cereal endosperm, and cassava roots, have plastids specialised for starch storage, called amyloplasts. Here, starch is stored for longer periods and is utilised in processes like sprouting and germination (Smith and Zeeman, 2020). Starch is composed of glucose units connected through linear α-1,4- and branch α-1,6-bonds. These form the highly branched amylopectin and the virtually unbranched amylose polymers (Apriyanto et al., 2022). Amylopectin chains form highly organised helices that pack tightly together, forming crystalline lamellae, while amylose chains are proposed to sit in amorphous lamellae between them. This gives starch granules a semi-crystalline, insoluble structure (Seung, 2020).

Although the semi-crystalline structure is highly conserved in plants, the number of starch granules per plastid and their individual morphology can vary greatly. In Arabidopsis chloroplasts, around five granules will form in each chloroplast, and they occur with flattened morphology in stromal pockets between the thylakoid membranes (Crumpton Taylor et al., 2012; Burgy et al., 2021). In wheat endosperm, one large, discoid A-type granule is formed per amyloplast, with multiple smaller, spherical B-type granules in stromules (Kamble et al., 2023). In rice endosperm, multiple granules are initiated within each amyloplast. As they grow, the granules impact each other and form compound granules, which have polyhedral shapes from their contact points (Matsushima et al., 2015). Potato has some of the largest known granules, with one ellipsoidal granule initiated per amyloplast (Hochmuth et al., 2025).

Although we know when and where granules are initiated in various species, relatively little is known about the mechanism of granule initiation. A critical step is the elongation of short, soluble maltooligosaccharides (MOS; Nakamura 2015; Seung and Smith 2019; Mérida and Fettke 2021; Kamble et al., 2023). There are many enzymes capable of extending these glucans in the amyloplast. These include starch synthases (SSs), a family of glucosyltransferases that extend linear chains by adding glucose units via new α-1,4-bonds, using ADP-glucose as a donor. SS1, SS2, and SS3 are involved in amylopectin synthesis, while Granule Bound Starch Synthase (GBSS) is involved in amylose synthesis (Delvallé et al., 2005; Zhang et al., 2008; Seung, 2020). In contrast, both SS4 and SS5 play an important role in granule initiation. Arabidopsis mutants lacking SS4 have only one or two granules per chloroplast compared with five to seven in wild type (Roldán et al., 2007; Crumpton-Taylor et al., 2013; Seung et al., 2016; Malinova et al., 2017; Lu et al., 2018). The granules in *ss4* mutants are initiated by SS3, as shown by the complete lack of starch granules in the *ss3 ss4* double mutant (Szydlowski et al., 2009). SS4 also plays an important role in granule initiation in amyloplasts, where wheat mutants lacking SS4 produced an increased number of initiations per amyloplast, resulting in compound granules (Hawkins et al., 2021). SS5 is closely related to SS4, but lacks catalytic glycosyltransferase activity. Despite this, Arabidopsis *ss5* mutants have fewer, larger granules per chloroplast, similar to *ss4* plants (Abt et al., 2020).

As well as starch synthases, α-1,4-linked glucose polymers can be extended by plastidial phosphorylase (PHS1), which uses the substrate glucose-1-phosphate, and is inhibited by ADP-glucose (Hwang et al., 2010). While in Arabidopsis *phs1* mutants lack a major starch phenotype, its absence exacerbates the granule initiation phenotype in *mex1* or *dpe2* double mutant plants (Malinova et al., 2013). In wheat endosperm, normal B-type granule initiation requires PHS1, while A-type granule initiation does not (Kamble et al., 2023). The potato PHS1 (called Pho1) has two known copies, designated as Pho1a and Pho1b, where Pho1a is more highly expressed in tubers (Sharma et al., 2023).

In this study, we investigated the role of SS4, SS5, and Pho1a in starch granule initiation by generating a set of CRISPR mutants with each gene individually knocked out. Subsequent analysis of the starch showed compound granules and supernumerary initiations in *pho1a* mutants, whereas granule morphology was not altered in *ss4* or *ss5* mutants. We propose a central role of Pho1a in starch granule initiation in tubers.

## Materials and Methods

### Plant Transformations

Potato (*Solanum tuberosum* L. *cv*. Clearwater Russet) was transformed using CRISPR-Cas9 ribonucleoproteins targeting either Pho1a (Soltu.DM.03G007710/Soltu.DM.03G007720 and Soltu.DM.03G007750/Soltu.DM.03G007760), SS4 (Soltu.DM.02G014060), or SS5 (Soltu.DM.02G027020). Targets within genes are indicated in **Table S1**. Clearwater Russet leaf protoplasts were isolated from tissue culture plantlets, transfected with CRISPR-Cas9 ribonucleoprotein (RNP) complexes, and subsequently regenerated as described in Andersson et al. (2018). RNP complexes targeting SS4, SS5, or PHO1a were assembled using the TrueCut Cas9 Protein v2 (Invitrogen) and the Alt-R™ CRISPR-Cas9 system (IDT). RNP transfection was performed by incubation with 250,000 protoplasts and 12.5% (w/v) PEG 4000 (Sigma-Aldrich) for 10 minutes. Leaf tissue was collected from regenerated tissue culture plantlets for genotyping by amplicon sequencing.

Gene-edited lines were genotyped using next generation sequencing. PCR amplicons were generated using primers flanking the Cas9 target sites (**Table S2**). Amplicons were sequenced by Illumina sequencing (MiSeq Nano, 2x250 bp), resulting sequences were clustered, and consensus sequences were aligned to the wild type sequences. Characterized edits are reported for *ss4* and *ss5* mutants in **Table S3 and S4**. *Pho1a* CRISPR-Cas9 mutants were subjected to proteomic analysis to confirm the absence of Pho1a (**Data 1**). Soluble proteins were extracted using 1 mL extraction solution (50 mM Tris-HCl, pH 8, 1 mM DTT, 1% [v/v] Triton X-100, 150 mM NaCl) for 200 mg freeze-dried tuber material at 4 °C. Insoluble material was removed by centrifugation. The proteins in the supernatant were precipitated using chloroform and methanol (Pankow et al. 2016). Peptides were identified and quantified using a nano-LC-MS/MS on an Orbitrap Eclipse Tribrid mass spectrometer with a FAIMS Pro Duo source, coupled to an UltiMate 3000 RSLCnano LC system (Thermo Fisher Scientific, Hemel Hempstead) according to Kamble et al. (2023).

Pho1a RNAi lines (named *siPho1a* to distinguish them from their CRISPR-Cas9-edited counterparts) were generated by *Agrobacterium*-mediated transformation. A binary vector pSIM5974 was cloned containing an inverted repeat targeting Pho1a 5’ UTR and coding sequence (NM_001288286 position 1-301). Vector construction and plant transformation was performed as described in Hochmuth et al. (2025). Briefly, Pho1a fragments were amplified from Clearwater Russet cDNA and confirmed by Sanger sequencing. The inverted repeat sequence was placed downstream the Cauliflower mosaic virus (CaMV) 35S promoter and upstream the ubiquitin-3 (UBI3) terminator. Clearwater Russet *siPho1a* lines were selected by PCR, using primers specific to the T-DNA insertion.

### Tuber phenotyping and determination of starch content

CRISPR-Cas9 edited plants were grown in glasshouse conditions with temperature control (18 °C minimum/27°C maximum) and supplemental lighting (16 h photoperiod with an intensity of about 1500 µmol/m^2^/s) (Simplot Plant Sciences, Boise, Idaho, USA). Plants were grown in 2-gallon pots for three months, where mature tubers were harvested. Pho1a-deficient tubers were photographed before processing. Tubers were weighed, lyophilised, and ground using metal beads and a Geno-Grinder (SPEX). Total and resistant starch content were determined using the Resistant Starch Assay Kit (Megazyme) as described in Navarro et al. (2026).

### Starch purification and analysis of granule characteristics

Starch purification was carried out on freeze-dried and ground potato tuber material. Approximately 300 mg of tuber powder was incubated in 5 mL NaCl (0.5 M) on ice for 30 min. The suspensions were passed through a 100 µm nylon mesh with an additional 5 mL of water. The starch was pelleted then resuspended in a 90% (v/v) Percoll solution (50 mM Tris-HCl, pH 8), and spun at 2,500*g* for 10 mins. The starch was washed in 2% (w/v) SDS three times, followed by three washes in cold acetone.

Granule size distributions were quantified using Multisizer 4e Coulter counter (Beckman Coulter, Indianapolis) fitted with a 200 µm aperture. Purified starch was suspended in Isoton II (Beckman Coulter) and a minimum of 20,000 particles were analysed per replicate.

Amylose content was estimated through iodine colorimetry (Knutson 1986). Purified starch (5 mg) was suspended in 10 mL of 6 mM iodine in DMSO, and rotated overnight to dissolve. A calibration curve containing five points ranging from 1-5 mg amylose (from potato starch, Sigma Aldrich) was included. This solution was diluted 1:10 in water and the absorbance at 600 nm was recorded using a SpectraMax 340PC spectrophotometer (Molecular Devices, California). The apparent amylose was calculated from the standard curve, and the amylose content calculated based on Knutson (1986).

Chain length distribution of purified starch granules was analysed using HPAEC-PAD as described in Navarro et al. (2026). Chain lengths were grouped for statistical analysis according to Hanashiro, Abe, and Hizukuri (1996) into very short (<6 degrees of polymerisation [dp]), A (6-12 dp), B1 (13-24 dp), B2 (25-36 dp), and B3 (>37 dp) chains.

### Microscopy

Purified starch was visualised using a Nova NanoSEM 450 (FEI, Hillsboro) scanning electron microscope. Before imaging, samples were coated with platinum (4-6 nm) using a ACE600 sputter coater (Leica, Newcastle upon Tyne). Light micrographs were captured using a Axio Imager Z2 (Zeiss, Oberkochen) fit with a polariser on purified starch stained with Lugol’s iodine solution.

To investigate localisation, we cloned Pho1a and LESV into plant expression vectors. RNA was extracted from developing Clearwater Russet tubers using the RNeasy kit (Qiagen), followed by GoScript Reverse Transcription System (Promega), according to manufacturers’ instructions. Pho1a coding sequence was amplified from total potato tuber cDNA, using the forward primer 5’-CACCATGGCGACTGCAAATGGAGC-3’ including the CACC overhang and the reverse primer 5’-TGCTATTTCCACAGCTTCAATGTTCCA-3’. The PCR product was cloned into the Gateway compatible pENTR vector using the pENTR/D-TOPO kit (Thermo), yielding Pho1a:pENTR. The Pho1a coding sequence was recombined into the Gateway-compatible plant expression vector, pB7RWG2 using LR Clonase II (Thermo), in frame with the C-terminal RFP tag. For LESV, the coding sequence from the LESV:pDONR221 entry vector (Locquet et al., 2026) was recombined into pB7YWG2, in frame with its C-terminal YFP tag. Three days after infiltration, leaf disks were imaged using a Zeiss LSM980 with Airyscan using a 63X water immersion lens. YFP was excited using light at 524 nm, RFP at 561 nm, and chlorophyll at 639 nm.

### Pull-down assay and Mass Spectrometry

To identify proteins that associate with Pho1a, bait protein (recombinant His-tagged Pho1a) was expressed in the *Escherichia coli* strain LOBSTR. The Pho1a coding sequence was recombined from the Pho1a:pENTR vector into the vector pTLV550 using LR clonase II. A control bait was prepared by transforming LOBSTR with an empty pProEXHtb vector (Invitrogen). *E. coli* were grown to an OD between 0.60 and 0.75 before adding 5 mM IPTG (Formedium) and incubating overnight at 18 °C. Cells were pelleted (50,000*g* 4°C for 10 min) the pellet was snap frozen and thawed on ice with 30 mL lysis buffer (50 mM TrisHCl, 300 mM NaCl, 40 mM Imidazole, 2 mM DTT, 1x cOmplete protease inhibitor cocktail [PiC; Roche]) per litre of *E. coli* grown. Once the pellet has mixed into the lysis buffer, the cells were sonicated on ice for 3 mins, followed by incubation at 4 °C for one hour. The lysate was separated from the pellet by centrifugation.

A protein extract was prepared from mature tubers from wild-type Clearwater Russet by homogenising tissue in ice-cold extraction medium (50 mM Tris-HCl, pH 8, 1 mM DTT, 1% [v/v] Triton X-100, 150 mM NaCl). Insoluble materials were removed by centrifugation. This soluble protein extract was mixed with the *E. coli* lysate containing the bait, and co-incubated with µMACS magnetic beads conjugated to anti-His for 1 hour at 4 °C (Miltenyi Biotec). The beads were captured using a magnetic stand (Miltenyi Biotec), washed and eluted according to Kamble et al. (2023). Peptides were identified and quantified using proteomic analysis (described above). Abundance ratios and p-values were calculated relative to the empty vector control.

### Yeast 2-hybrid assays

For Yeast 2-hybrid (Y2H) assays, coding sequences from Pho1a:pENTR, LESV:pDONR221, and ISA1:pDONR221 (Locquet et al. 2026) were recombined directly into Gateway-compatible pGADT7 (in frame with N-terminal Activation Domain) or pGBKT7 (in frame with N-terminal Binding domain). These constructs were transformed into yeast using the EZ-Yeast transformation kit (MP), alongside pBD-53 + pAD-T as a positive control, and assays were conducted as described in Chen et al. (2024).

### Native gel visualisation of phosphorylase activity

Pho activity was measured on mature, glasshouse grown Clearwater Russet and Desiree tubers, based on Zeeman et al. (2004) with minor modifications. Soluble proteins were extracted from tuber material using extraction buffer (100 mM DTT, 1 mM MOPS, pH 7.2, 1 mM EDTA, 10% [v/v] ethanediol, and 1x PiC) and spun at 20,000*g* for 10 min to remove insoluble material. Protein concentration in the supernatant was quantified using Bradford Assay. Equal amounts of protein were added to each well of a native PAGE gel (using 7.5% acrylamide and 0.3% oyster glycogen in the resolving gel, and 3.75% acrylamide in stacking gel), followed by electrophoration at 4°C at 100V for 4 hours. The native PAGE gel was washed twice in wash buffer (100 mM Tris-HCl, pH 7.0, and 1 mM DTT) and incubated overnight in Glucose-1-Phosphae solution (100 mM Tris-HCl, pH 7.0, 1 mM DTT, 50 mM Glc-1-P) at room temperature. Activity bands were visualised by staining with Lugol’s solution (Sigma, St. Louis).

### Statistical analyses

One-way ANOVA with Tukey’s Honest Significant Difference (HSD) post-hoc test was used for mean granule size, starch content, amylose content, tuber number, tuber mass, and dry matter content. For chain length distribution, pairwise t-tests were used with Bonferroni correction applied to the p-value. Analyses were carried out using R version 4.2.1.

## Results

### Loss of Pho1a, but not SS4 or SS5, affects starch granule phenotypes

Knockout mutants lacking one of the glucosyltransferases implicated in starch granule initiation, Pho1a, SS4, or SS5, were generated using CRISPR-Cas9. Thirty *ss4* lines and twenty-three *ss5* lines were generated. The lines were genotyped using amplicon sequencing over the edit site, and five *ss4* lines and four *ss5* lines with no detectable wild type alleles were selected for further investigation (**Tables S2 and S3**). Henceforth, *ss4* and *ss5* lines refer to CRISPR-edited lines with no detectable wild type alleles (i.e. full knockouts). Fifteen *pho1a* lines were generated, but genotyping by amplicon sequencing was inconclusive due to a complex genotype at Pho1a, with at least two and up to four copies per chromosome, giving up to 16 copies in the tetraploid genome (Sharma et al., 2023). We therefore kept all lines for phenotypic screening.

Starch granules were extracted from mature tubers for examination of granule morphology. Granules from wild-type potato tubers were large and ellipsoidal (**Figure 1A**,**C**). When viewed under polarised light, a single maltese cross was visible, indicating a single centre of organisation (initiation point) per granule (**Figure 1B**). Starch from *ss4* and *ss5* tubers showed no visible morphological differences to the wild type, and also showed a single initiation point within each granule. We screened all *pho1a* lines for changes in granule size, and every line had smaller starch granules than the WT, so we selected the five lines with the smallest average granule size for further analysis (numbered line 1-5) (**Figure S1**). These *pho1a* lines showed distinct differences to the wild-type starch granules. The granules were smaller and showed varying morphologies including ellipsoidal (as in the wild type), polygonal, and irregular (**Figure 1A, C**). Under polarised light microscopy, the ellipsoidal and polygonal granules contained single initiation points, while others were compound granules, formed of many small granules, giving irregular shapes with a range of levels of surface confluence. To confirm that these phenotypes were associated with the absence of Pho1a, we used proteomics, and no peptides corresponding to Pho1a were detected in tuber protein extracts (**Table S5**). These five lines with no detectable Pho1a protein expression are henceforth referred to as *pho1a* lines.

**Figure 1:**
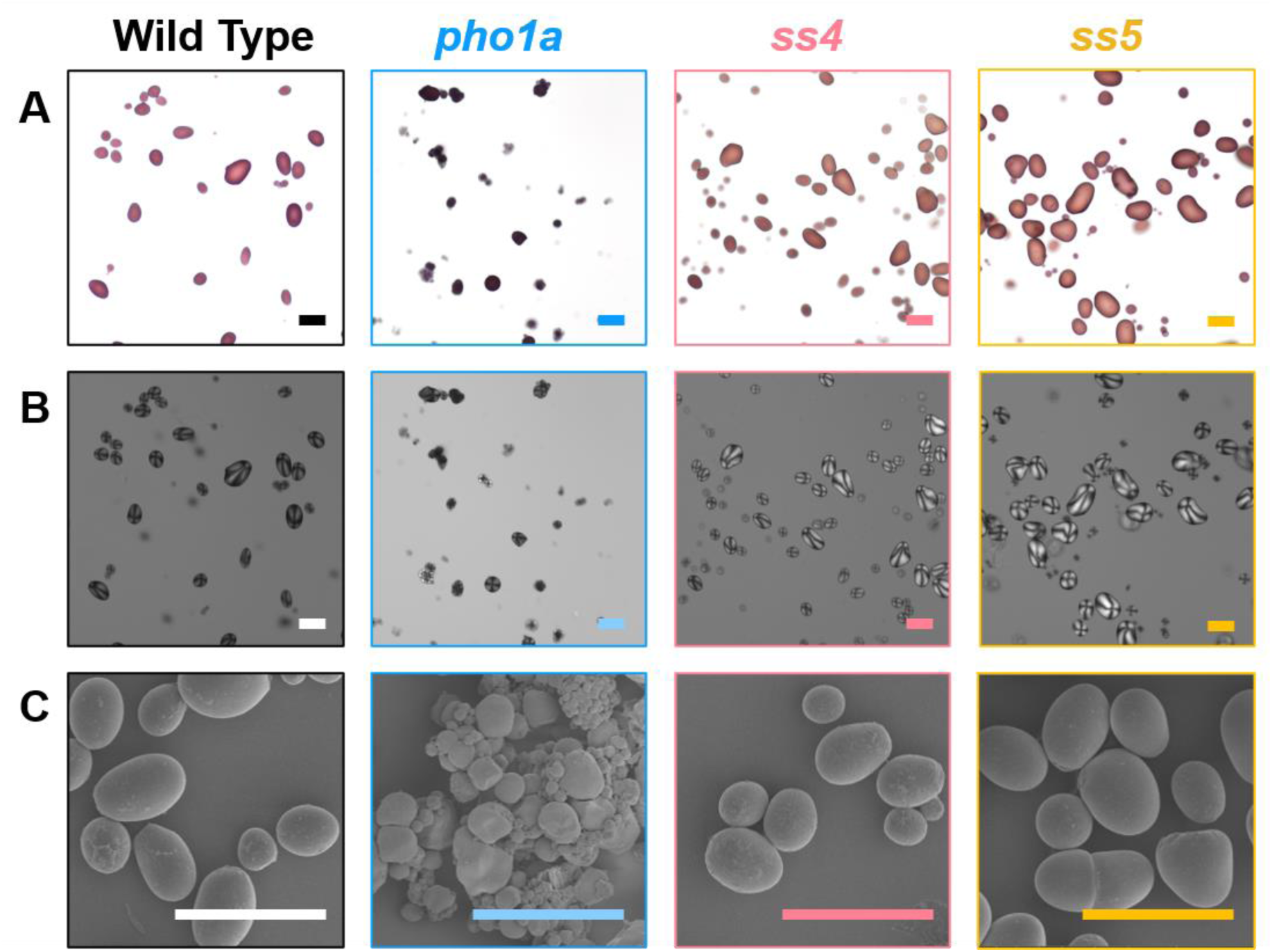
Granule morphology in pho1a, ss4, and ss5 in purified tuber starch. Scale bars = 50 µm. **A)** Brightfield images of iodine-stained starch granules. **B)** Images as in A, with polarised light. Representative images are shown from the lines pho1a-3, ss4-3, and ss5-4. **C)** Scanning electron micrographs of extracted starch. Representative images are shown from the lines pho1a-4, ss4-1, and ss5-2.

We next analysed granule size and starch structure in the lines. Quantification of granule size using the Coulter counter showed that *ss4* and *ss5* starch had no change in granule size (**Figure 2A**). There was also no change in amylose content in these mutants (**Figure 2B**), and the tubers had equivalent levels of total and resistant starch (i.e. not readily digested starch) compared to the wild type (**Figure 2C**). The *ss4* starch showed minor alterations in amylopectin chain length distribution, with slightly fewer A-chains (6-12 dp) compensatory increases in B1 (13-24 dp) and B3 (>36 dp) chains (**Figure 2D; Figure S2A**). In *pho1a* lines, granule size was almost half that of the wild type, with markedly less resistant starch in each line (**Figure 2A**,**C**). While no change was observed in the amylose content, the chain length distribution of *pho1a* starch differed significantly from the wild type (**Figure 2B, D; Figure S2C**). The *pho1a* lines had significantly more very short (<6 dp), B2 (25-36 dp) and B3 chains. There was a compensatory decrease in B1 chains in *pho1a* lines, and no change in the A-chains. Of the mutants, *pho1a* were the only lines with significant alterations in starch content and digestibility, where resistant starch content was greatly reduced.

**Figure 2:**
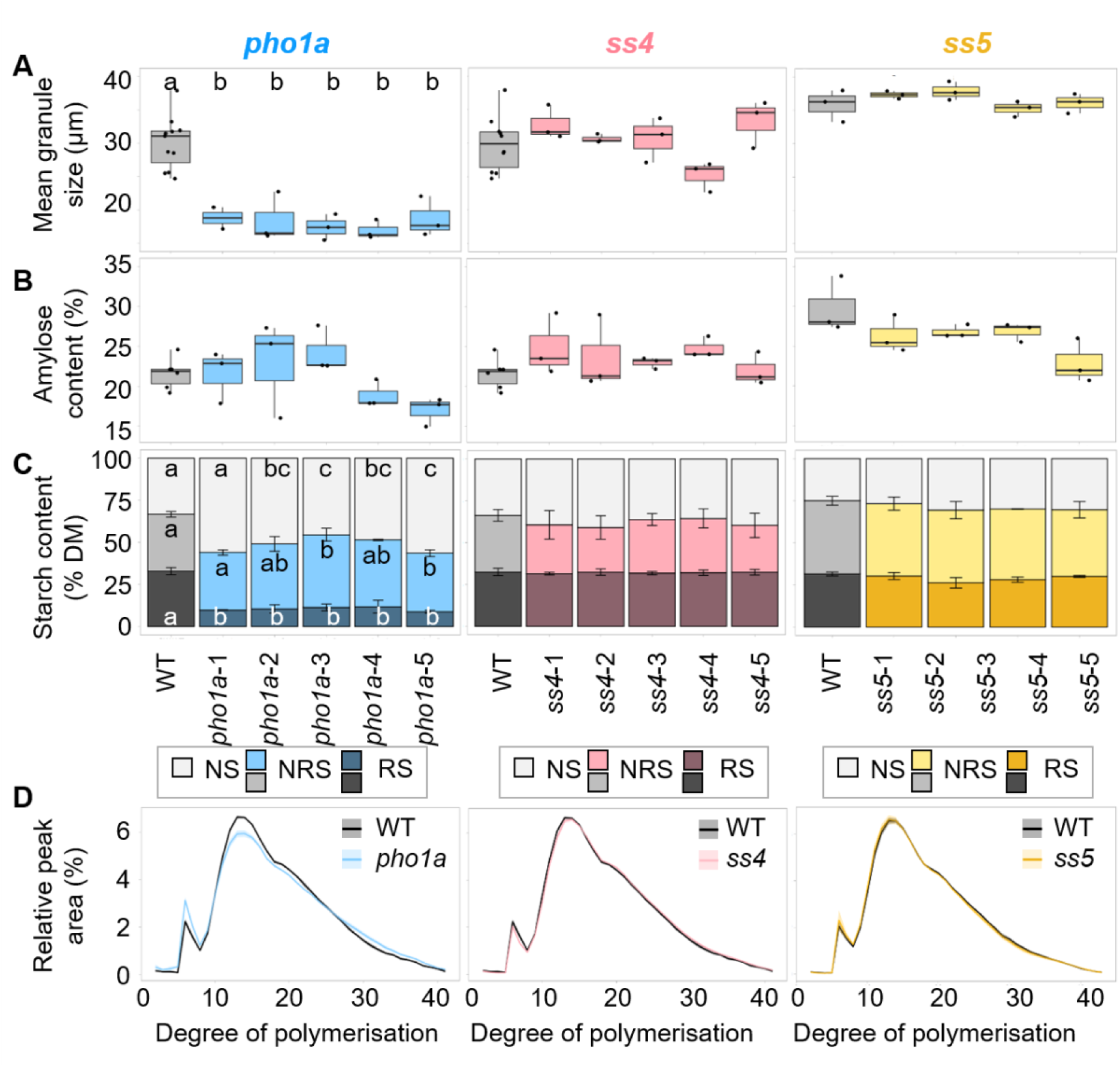
Starch granule characteristics from pho1a, ss4, and ss5 CRISPR mutant tubers. **A)** Granule size was quantified using a Coulter counter, and data are represented as the mean from each plant, n ≥ 3. **B)** Amylose content as determined through an iodine-binding assay. **C)** Resistant starch (RS), non-resistant (NRS), and non-starch (NS) content is represented as a percentage of the dry matter (DM). Error bars represent the standard deviation (SD) of the RS and NRS fractions, using three biological replicates per line. For A-C, lines with different letters are significantly different under a one-way ANOVA with Tukey’s Honest Significant Difference (HSD) post-hoc test (p <0.05). Panels with no lettering have no significant differences. **D)** Chain length distribution of amylopectin in 4 - 5 mutant lines versus the wild type (n ≥ 3 biological replicates per line). The solid line represents the mean, with shading ± standard deviation.

### RNAi *siPho1a* lines have a similar granule phenotype but no tuber phenotype

The *pho1a* plants had a distinct tuber phenotype. The tuber yield per plant was reduced by ~75% compared to wild type (**Figure 3A**). This was due to lower tuber mass, as the same number of tubers were harvested in the *pho1a lines* and the wild type (**Figure 3B, C**). Furthermore, tubers in *pho1a lines* had nearly a third lower dry matter than their wild-type counterparts, and an elongated shape with atypical protrusions (**Figure 3D**,**E**). Given the negative impacts on tuber yield and morphology, we generated RNAi lines that likely contained some residual Pho1a activity. From the 14 *siPho1a* lines generated, five with the lowest average granule size were selected for further analysis. In these *siPho1a* lines, similar granule characteristics were seen as in the *pho1a* lines, with a ~40% reduction in average granule size compared to the WT and the formation of compound and polygonal granules (**Figure 4**). Notably, the distinctive tuber phenotype observed in the CRISPR *pho1a* lines was not seen in the RNAi *siPho1a* lines, despite the similarities in the granule phenotype. No significant reduction in tuber mass was seen in any *siPho1a* line, and while two of the five lines had reduced dry matter content, the overall reduction was only ~13%, less than half of the value for the CRISPR lines. Most notably, the tuber morphology was nearly indistinguishable from the wild type, with only a slight elongation visible (**Figure 3E**).

**Figure 3:**
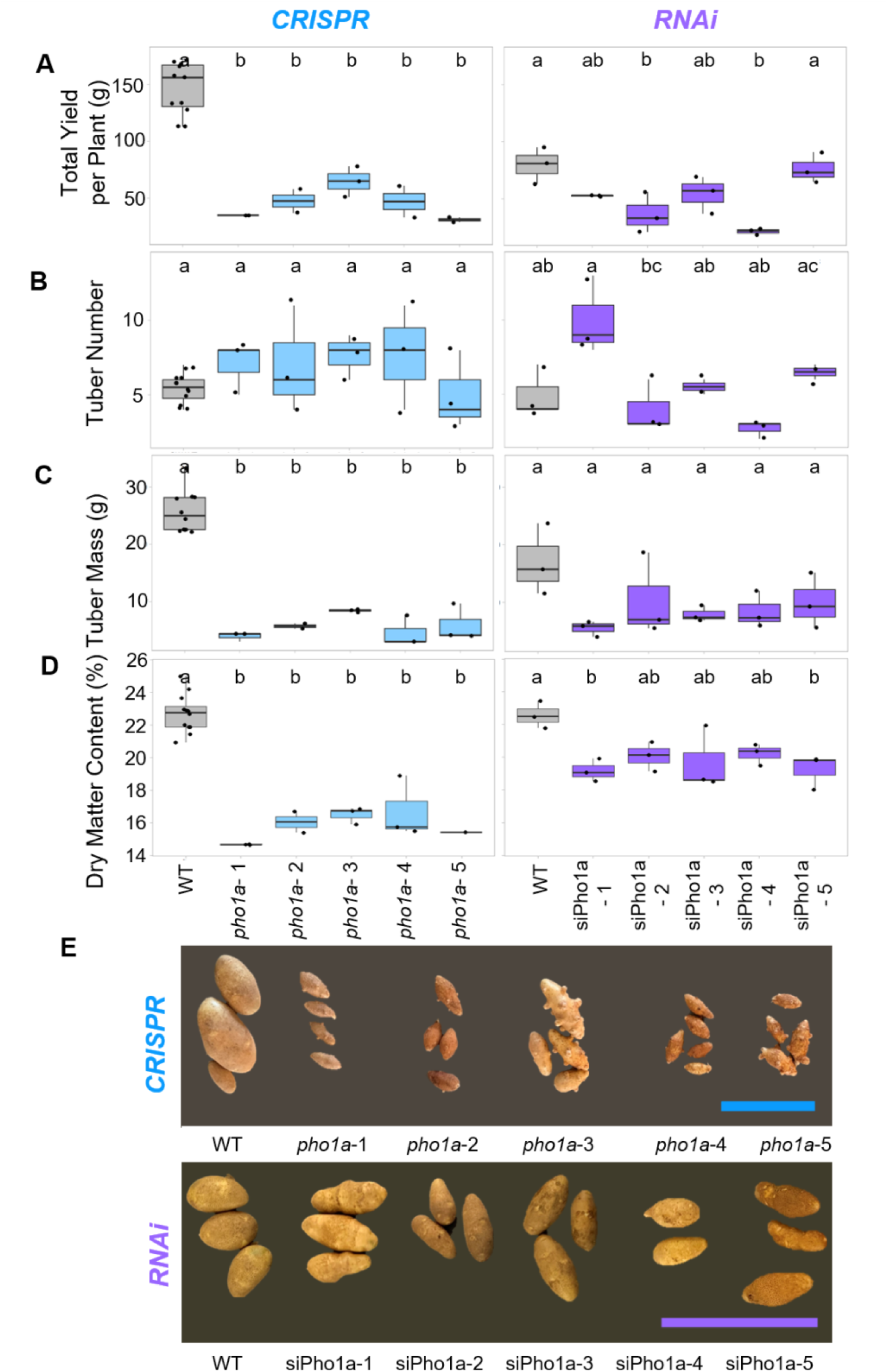
Tuber phenotypes in Pho1a-deficient plants in CRISPR and RNAi lines. **A)** Total yield was measured as weight of all tubers produced by a single plant. **B)** Number of tubers produced per plant. **C**) Average mass of individual tubers, calculated by dividing the total yield by the number of tubers per plant. **D)** Dry matter content in Pho1a-deficient plants. For A-D, lines with different letters are significantly different under a one-way ANOVA with Tukey’s Honest Significant Difference (HSD) post-hoc test (p <0.05), n ≥ 3 biological replicates per line. **E)** Tubers from Pho1a-deficient plants. Scale bar = 10 cm.

**Figure 4:**
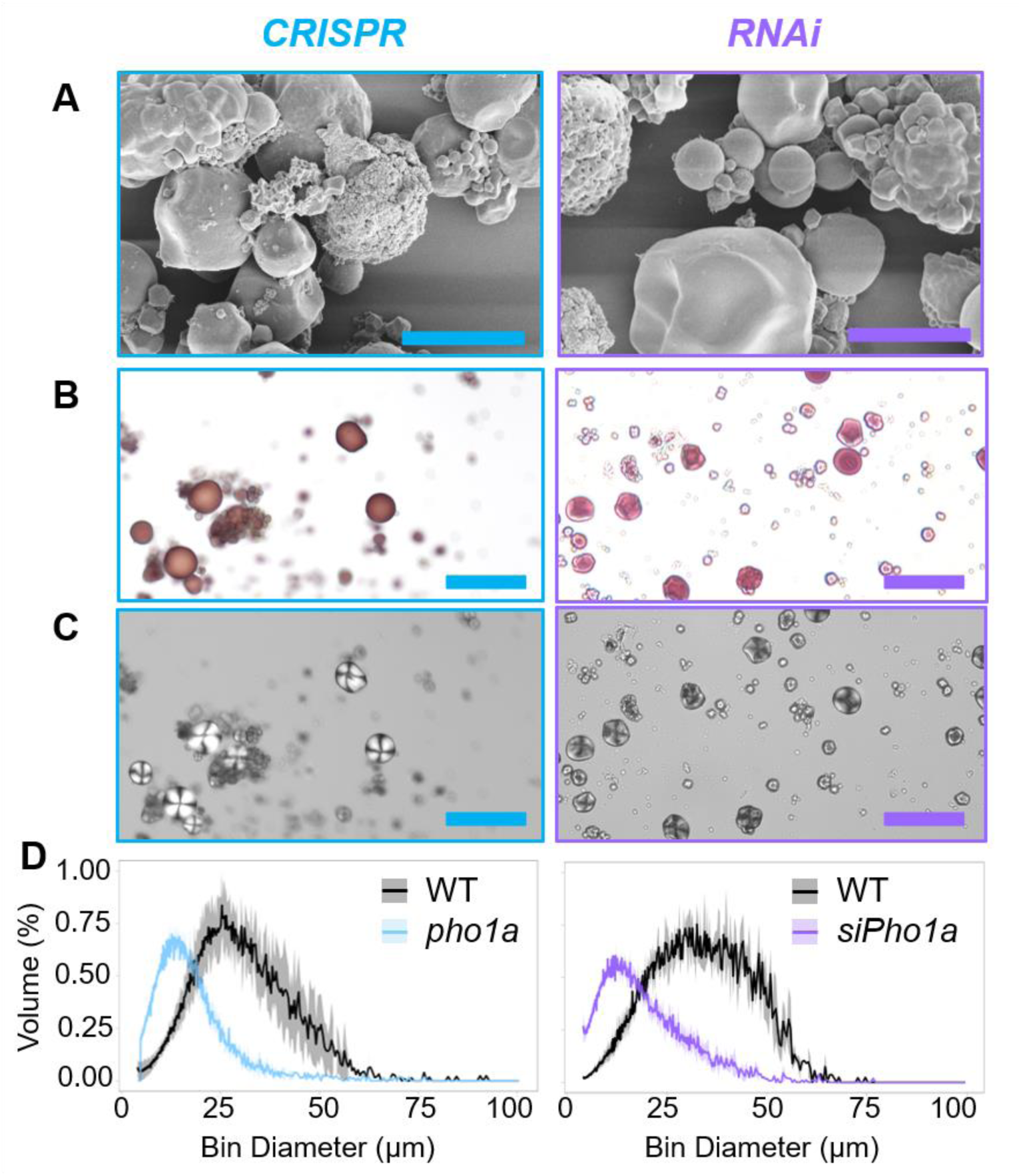
Starch granule phenotypes in tubers deficient in Pho1a. CRISPR lines (blue) are full knockouts for Pho1a, and purple represents RNAi siPho1a lines. **A)** Scanning electron micrographs of extracted starch. Scale bars = 10 µm. Representative images are shown from the lines pho1a-3 and siPho1a-5. **B)** Bright-field and **C)** polarised light images of extracted tuber starch. Representative images are shown from the lines pho1a-3 and siPho1a-1. Scale bars = 50 µm. **D)** Granule size distribution of extracted starch quantified on the Coulter Counter. Solid lines represent the mean from n ≥3 replicates for the wild type, and 5 individual lines containing 3 biological replicates for pho1a and siPho1a lines, and the shading represents the standard deviation.

Given the differences between our findings and the *pho1a* mutants generated in Solanum tuberosum cv. Desiree by Sharma et al. (2023), we investigated the differences between Pho activity in the two cultivars. Visualising phosphorylase activity on native PAGE gels showed that Clearwater Russet tubers had stronger bands compared to Desiree, indicating more Pho1 activity in Clearwater Russet (**Figure S3**).

### Pho1a and LESV colocalise to the starch granule

Based on the importance of Pho1a in granule initiation, we investigated proteins that were associated with Pho1a using affinity pull-down and MS. Pho1a with an N-terminal His-tag (His-Pho1a) was expressed in *E. coli*, and the bacterial lysate was co-incubated with a soluble protein extract from mature Clearwater Russet tubers. Anti-His beads were used to pull-down His-Pho1a and associated proteins, which were subsequently identified and quantified using mass spectrometry. Over a thousand plant proteins were enriched at least 2-fold (**Data S1**), and those that appeared in our list of starch metabolic proteins in potato (**Table 4**) were shortlisted, and these 19 shortlisted proteins are reported in **Table 1**. We focused on LIKE EARLY STARVATION (LESV), because it was among the most enriched proteins in the affinity pull-down, and was previously reported to have a similar small-granule phenotype as *pho1a* when knocked out (Locquet et al., 2026).

**Table 1:**
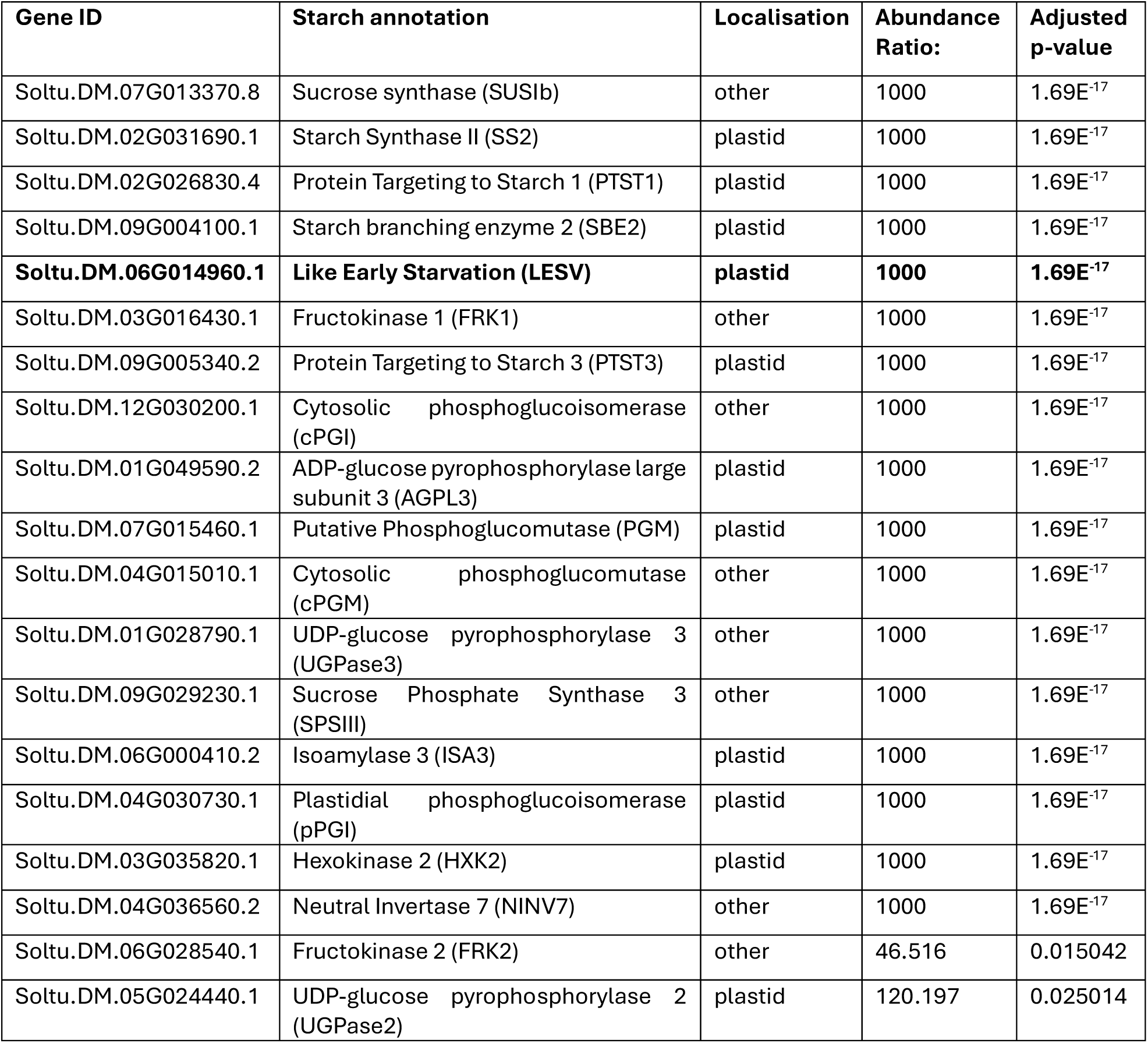
Starch metabolic proteins significantly enriched by the Pho1a pull-down. Abundance ratio was calculated using the abundance of each protein in the Pho1a pull-down vs the control. A ratio of 1000 represents the maximum enrichment value, indicating the protein was detected in the Pho1a pull-down but not in the control. The adjusted p-value indicates statistical significance of the difference in protein abundance between the Pho1a pull-down and the control.

Based on these findings, we conducted yeast 2-hybrid (Y2H) assays to confirm whether the two proteins interact directly. The co-transformation of LESV and Pho1a did not result in colony growth, suggesting that the two proteins do not directly bind each other, or cannot interact within the yeast nucleus (**Figure 5A; Figure S4**). Notably, Pho1a also did not interact with ISA1, which is known to interact with potato LESV and produce similar phenotypes when suppressed. Given prior evidence that LESV can facilitate ISA1 localisation to starch granules in rice (Yan et al. 2024), we investigated whether LESV could alter localisation of Pho1a, using transient expression of fluorescently-tagged proteins in *N. benthamiana* leaves. Both LESV-YFP and Pho1a-RFP proteins located in the plastid. When LESV-YFP was expressed alone, it appeared around the surface of starch granules within chloroplasts, appearing ring-like in confocal laser-scanning micrographs (**Figure 5B**). By contrast, Pho1a-RFP when expressed alone showed a diffuse signal throughout the chloroplast, indicating stromal localisation. When both proteins were expressed together, Pho1a-RFP localisation shifted, appearing mainly on the granule surface together with LESV-YFP, with only a low level of stromal signal. This co-localisation suggests a possible functional interaction between Pho1a and LESV *in planta*.

**Figure 5:**
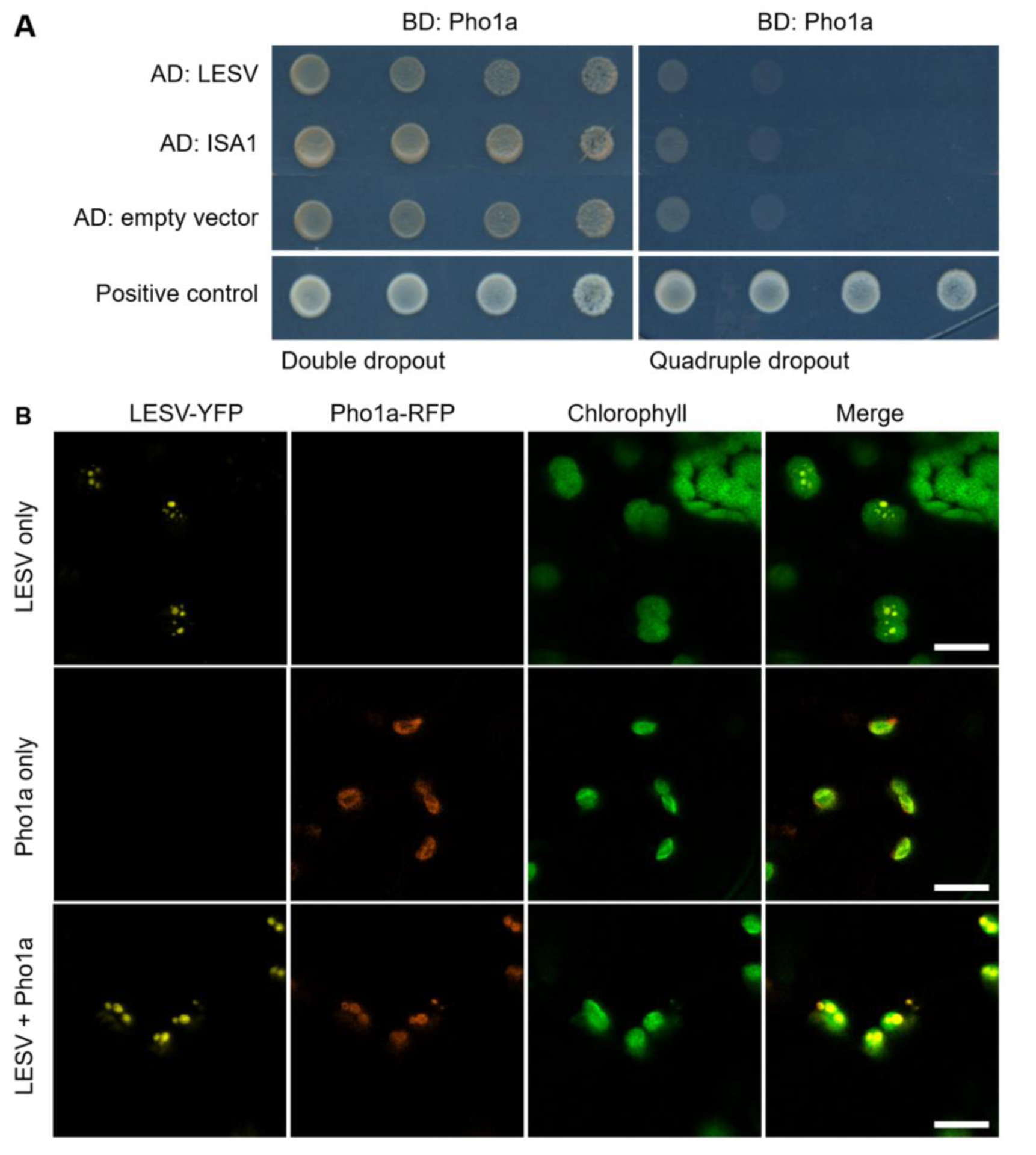
LESV alters the location of Pho1 in Nicotiana benthamiana chloroplasts. **A)** Yeast 2-Hybrid (Y2H) assays showing no interaction between Pho1a and either LESV or ISA1, with either the activation domain (AD) or binding domain (BD). Double dropout medium (-Leu-Trp) was used to show growth, while quadruple dropout medium (-Ade-Leu-Trp-His) was used to show physical interaction with Pho1a. Positive control represents BD-p53 + AD-T. **B)** Confocal laser-scanning micrographs showing the localisation of LESV and Pho1a in Nicotiana benthamiana leaves. LESV-YFP, Pho1a-RFP, or both were transiently expressed with 35S promoters. Scale bars = 10 µm.

## Discussion

### Granule initiation in potato tubers is driven by Pho1a

We investigated starch granule initiation in potato tubers by systematically analysing the role of glucosyltransferases implicated in starch granule initiation in other species. We discovered that granule initiation in this organ is mainly driven by Pho1a, rather than SS4. Pho1a-deficient Clearwater Russet tubers produced compound granules, with reduced size, polygonal and irregular morphologies, and increased susceptibility to digestion (**Figure 1**, **2**). In contrast, tubers lacking SS4 or SS5 showed no difference in granule size or morphology, with no changes in amylose or starch content.

Similar starch granule phenotypes to our *pho1* mutants were reported in potato tubers with reduced SS3 (Abel et al., 1996), ISOAMYLASE (ISA) 1 or ISA2 (Bustos et al., 2004), and LESV (Locquet et al., 2026) protein levels. This suggests that this suite of proteins is important for the control of starch granule initiation in potato. This differs from the suite of proteins defined in Arabidopsis leaves (Seung and Smith, 2019; Uauy et al. 2025). Arabidopsis single mutants lacking either SS3, LESV, or PHS1 (the Arabidopsis ortholog of Pho1a) are not reported to have altered starch granule number per chloroplast (Malinova et al., 2013; Malinova and Fettke, 2017). However, ISA appears to be important in controlling granule number across multiple species including Arabidopsis, barley, and potato (Delatte et al., 2005; Burton et al., 2002; Bustos et al., 2004). Within this suite of proteins, our study positions Pho1a alongside LESV as playing a central role in granule initiation in potato tubers. The affinity pull-down experiment from potato tuber extracts suggested that Pho1a and LESV interact (**Table 1**), while the Y2H did not show direct protein-protein interaction (**Figure 5**). It is possible that the interaction between the two proteins is indirect, such that the interaction is facilitated by another protein that binds both LESV and Pho1a or a shared glucan substrate. The absence of additional plant proteins and glucan substrates in the yeast nucleus could explain why the interaction only appears *in planta*. Consistent with a substrate-mediated interaction, LESV was able to facilitate the localisation of Pho1a to starch granules when co-expressed in *Nicotiana* chloroplasts. This is not the first time Pho1a and LESV activities have been linked: Singh et al (2022) found that Arabidopsis PHS1 had reduced activity when incubated with LESV. Based on the pull-down, co-localisation, and the similarities of phenotypes, there is strong evidence that Pho1a and LESV have a functional interaction that affects starch granule initiation.

Sharma et al. (2023) noted a similar, but less severe, starch phenotype in Pho1a knockout tubers in the potato cultivar Desiree. Their mutant had smaller granules than the wild type with multiple initiations per amyloplast, but did not observe compound granules like we did in our mutants in Clearwater Russet. Interestingly, Sharma et al. (2023) found that CRISPR mutants retaining at least one wild-type allele (i.e. retaining some Pho1a activity) developed normal granules in size and number, in contrast to our RNAi lines which had granules indistinguishable from the full CRISPR knockouts (**Figures 3** and **4**). There are several potential reasons for different results between our study and Sharma et al. (2023). Pho1a activity was higher in Clearwater Russet compared to Desiree, supported by the stronger bands seen in the native PAGE (**Figure S3**), which could reflect differences in G-1-P metabolism between the varieties, contributing to differences in phenotypic severity following Pho1a elimination. We cannot rule out additional factors, including the possibility of Pho1a sequence differences across the multiple copies and chromosomes, which could have led to some alleles not being edited or being incorrectly genotyped in the respective cultivars. There also may be differences in MOS availability between the cultivars. Pho1b is thought to be exclusively expressed in cells close to vasculature in tubers in Desiree (Albrecht et al., 2001), but varietal differences in this expression pattern is not known. Pho1b is highly similar in protein sequence to Pho1a and likely has similar functionality that could compensate in the absence of Pho1a.

The elimination of Pho1a resulted in supernumerary initiations in potato, rather than fewer starch granules. Although the exact mechanism underpinning this phenotype is not known, it is plausible that the loss of key granule initiation proteins in tubers results in uncontrolled, increased granule initiations. In the wild type, tuber amyloplasts natively contain a single starch granule, which could be initiated by Pho1a. The initiation of this granule likely provides the correct substrate for all other starch-synthesising proteins to further develop that granule. Therefore, the presence of the single initiation suppresses other initiations by sequestering enzymes to the growing granule. However, in the absence of a granule (by the removal of Pho1a), other glucosyltransferases may elongate non-preferred substrates such as MOS and initiate more granules. A similar explanation was proposed for the supernumerary initiations seen in the endosperm of wheat *ss4* mutants (Hawkins et al., 2021). Indeed, all starch synthase isoforms typically involved in amylopectin synthesis (SS1-3) can elongate MOS in vitro when provided with no other substrates (Brust et al. 2013). Further, Arabidopsis mutants lacking both main granule initiation enzymes, SS3 and SS4, can initiate starch granules following the additional elimination of starch hydrolytic activity (AMY3) – suggesting that other glycosyltransferases can initiate granules under the right conditions (Seung et al. 2016). There are therefore multiple glucosyltransferases in plants that are capable of initiating granules, allowing for diversification of roles in initiation and biosynthesis, both within and between species.

### Starch granules initiate and grow normally in Clearwater Russet tubers lacking SS4 and SS5

SS4 is a central player in starch granule initiation in all plant species examined thus far. However, our work discovered that SS4 is not required for normal granule initiation in potato tubers. We report no change in starch content, granule size, or granule morphology in the complete absence of SS4 (**Figures 1** and **2**). In contrast, Arabidopsis mutants lacking SS4 have a dramatic reduction in starch granule number, and major changes in starch content and granule morphology (Roldán et al., 2007). Likewise, in wheat leaves and endosperm, loss of SS4 has a major effect on starch granule number (Hawkins et al., 2021). In rice, there are two isoforms of SS4 (SS4a and SS4b), and knocking out SS4b alone reduced starch content and granule size in developing endosperm (Toyosawa et al., 2016). This phenotype was not as pronounced as in Arabidopsis or wheat, which could be due to the presence of SS4a. While we did not see any effect on starch granule size or morphology in the *ss4* mutant of potato, the mutant had minor changes in amylopectin chain length distribution, which was similar to changes seen in Arabidopsis mutants (Roldán et al., 2007). In contrast, there was no alteration in amylopectin chain length structure in wheat (Hawkins et al., 2021). This suggests that the role of SS4 varies substantially between species and organs, and in potato tubers it contributes to starch biosynthesis by extending amylopectin chains rather than in granule initiation. Further exploration of other species may provide more examples of roles of SS4 outside granule initiation. Bioinformatics and biochemical analysis revealed that granule initiation may occur independently of SS4 in green algae *Chlamydomonas reinhardtii*, which warrants further genetic exploration (Courseaux et al., 2025).

SS5-deficient Arabidopsis mutants formed fewer and larger starch granules in their chloroplasts (Abt et al., 2020). However, SS5 in Arabidopsis is a substantially different protein from the potato SS5, as the Arabidopsis protein has a truncated GT-1 domain. Hu et al. (2025) used CRISPR-Cas9 to target the 5’ untranslated region, knocking down SS5 in the dihaploid potato line AC142. In these tubers, up to six starch granules formed in each amyloplast, and the starch also had higher amylose content. This strongly contrasts the lack of changes seen in the Clearwater Russet ss5 mutants presented in this study (**Figure 1**, **Figure 2**). Dihaploids will intrinsically have less allelic diversity than tetraploid cultivars like Clearwater Russet, as well as unique metabolic and genetic backgrounds, just as we saw differences between Clearwater Russet and Desiree. Therefore, differences between our findings and Hu et al. (2025) could also be due to differences in the genetic backgrounds.

### Pho1a has multiple roles in starch synthesis

In addition to its role in starch granule initiation, we propose that Pho1a is also involved in granule biosynthesis through extension of amylopectin chains in potato amyloplasts, based on the observed differences in chain length distribution in the *pho1a* lines (**Figure 2D**). The increase in short chains and decrease in B1 chains is similar to that reported for the rice *pho1* mutants (Satoh et al. 2008). The *pho1a* lines also had a clear tuber morphology phenotype, with protrusions forming a knobby appearance. This has historically been categorised as secondary growth, where growth under normal conditions is interrupted by high temperature stress. However, the cellular or biochemical mechanisms for secondary growth remain greatly unknown (Lugt et al., 1964; Armstrong and Grice, 1988; Sonnewald and Sonnewald, 2014). While environmental conditions were not a factor tested in this study, it is possible that the complete loss of Pho1a caused stress during growth, or sensitised these tubers to heat stress. Notably, Russet potatoes are particularly susceptible to secondary growth (Armstrong and Grice, 1988), which may explain why Sharma et al. (2023) did not observe the same phenotype in their *pho1a* tubers.

Commercialisation of the *pho1a* lines is likely precluded by their poor yields and knobby tubers (**Figure 3**). However, since the starch phenotype in the RNAi lines appears identical to the CRISPR lines, without the reduction in tuber size or appearance of protrusions, the starch phenotype may have commercial value. We noted an increase in digestibility of the raw starch by α-amylase, which was likely due to the increased surface area of the smaller and compound granules. However, the differences in chain length distribution may also have contributed to this increase in digestibility, or the additional surface area may have led to differences in surface modifications. Although not quantified in this study, there is also possibly deviations in the crystallinity and phosphorylation levels of the starch, which may have also affected the digestibility. The low volume of tubers generated in this study precluded assessment of the starch rheological properties, but would be a valuable direction of future study and guide potential utilisation of this phenotype in food products or industrial applications. The expanding knowledge of proteins are involved in starch granule initiation and biosynthesis in potato tubers gives valuable insight for breeders, particularly given the striking differences between potato and other species.

## Acknowledgements

We acknowledge JIC Bioimaging platform for providing access to microscopes, and the Proteomics platform for sample analysis. This work was funded through a Biotechnology and Biological Sciences Research Council (BBSRC, UK) Industrial Partnership Award BB/X001520/1 (to D.S.), the BBSRC-funded Institute Strategic Programme Harnessing Biosynthesis for Sustainable Food and Health (HBio)(BB/X01097X/1), and the Deutsche Forschungsgemeinschaft (DFG FE 1030/ 5-1, 6-1 and CRC 1644 to J.F.).

