## Supplementary Figures and Tables for "Plastidial Phosphorylase (Pho1a) is the dominant glucosyltransferase regulating starch granule initiation in potato tubers"

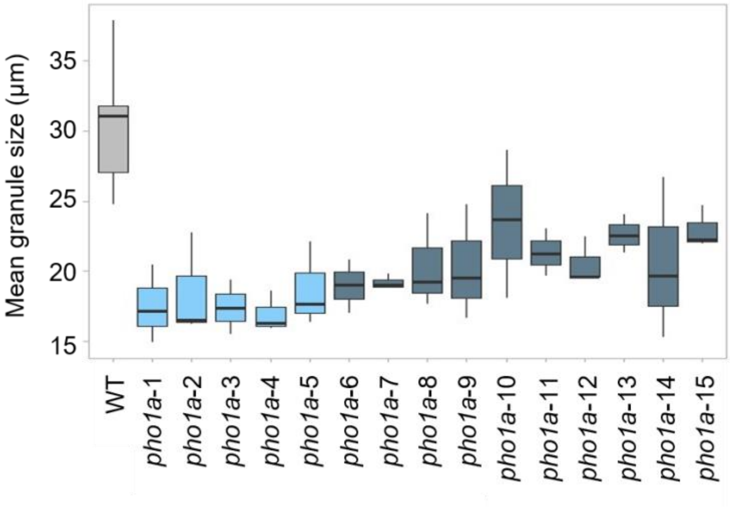


***Figure S1: Average granule size in tubers from independent pho1a lines.*** *Average size was calculated from Coulter counter data. Selected lines are represented in light blue and labelled corresponding to their naming in Figure 2.*

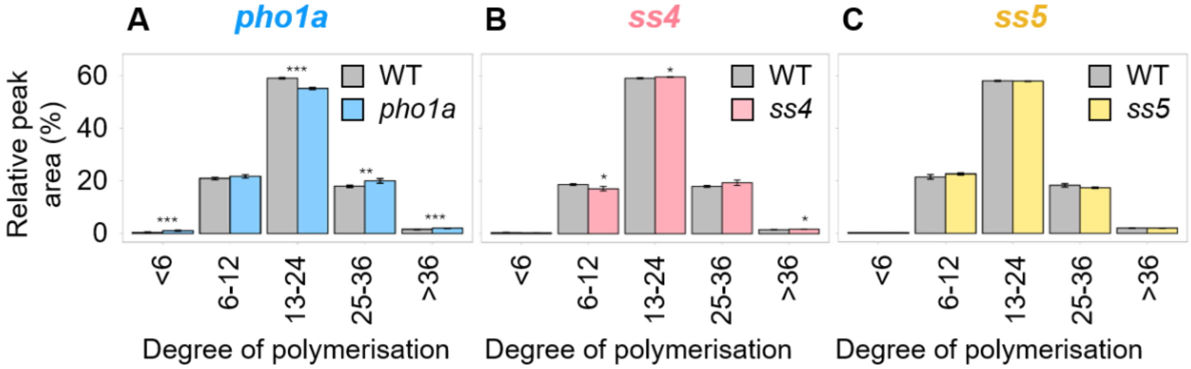


**Figure S2: Chain length distribution of amylopectin in A) *pho1a*, B) *ss4*, and C) *ss5* lines.** Amylopectin chain length distribution in 4 - 5 mutant lines versus the wild type, *n ≥ 3 biological replicates per line*. The chains are grouped according to Hanashiro, Abe, and Hizukuri (1996). Significant differences to the wild type based on pairwise t-tests are represented by * ($p < 0.05$), ** ($p < 0.01$), or *** ($p < 0.001$).

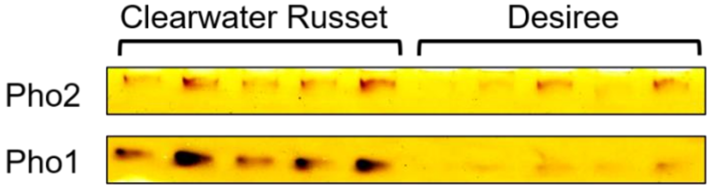


**Figure S3: Native PAGE of phosphorylase activity in Clearwater and Desiree tubers.** Gels were loaded on an equal protein basis, with 100 µg loaded in each lane.

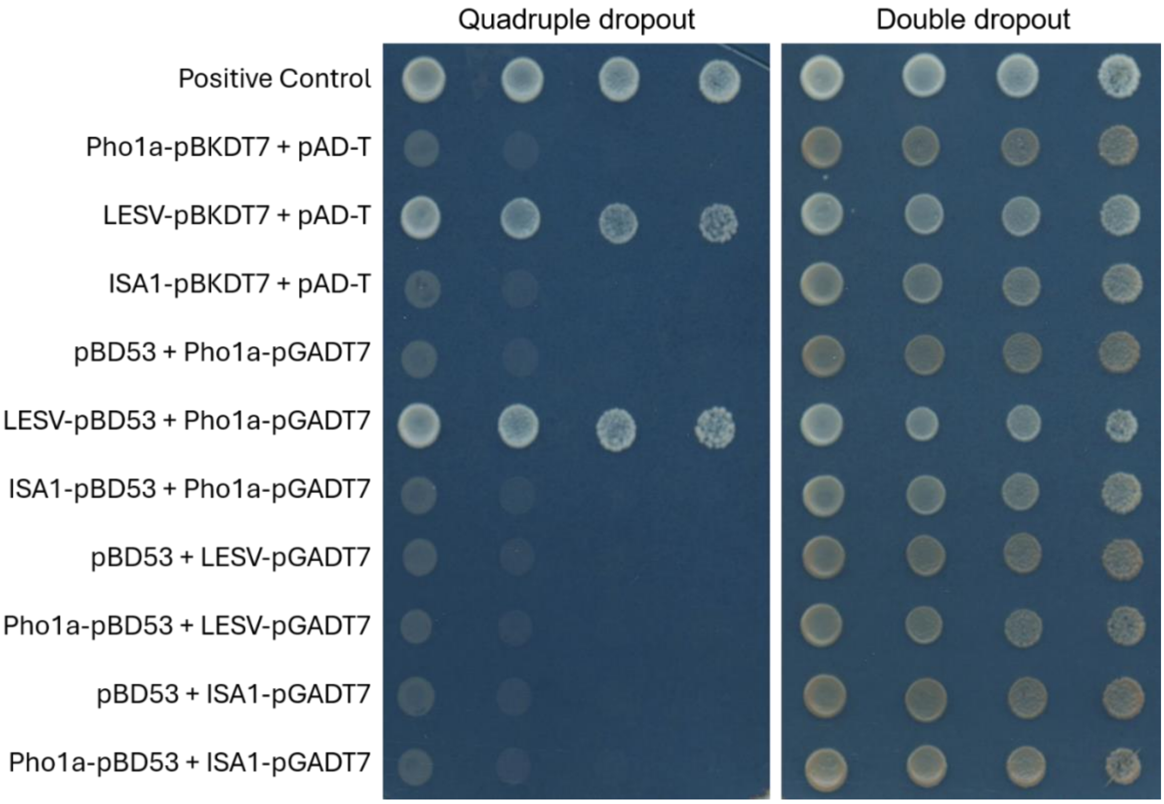


**Figure S4: Yeast 2-Hybrid (Y2H) assays.** Double dropout medium (-Leu-Trp) was used to show successful transformation of plasmids, while quadruple dropout medium (-Ade-Leu-Trp-His) was used to show protein interactions.

**Table S1. CRISPR Cas9 gRNA Sequences.**

| **Gene** | **Target Sequence (Top Strand)** | **Target location** | **gRNA Sequence (5' to 3')** |
|---|---|---|---|
| SS4 | CCATATTGCAGCCGAGATGGCGC | Exon 5 | GCGCCATCTCGGCTGCAATA |
| SS5 | TTGTACTGAAATGGCACCAGTGG | Exon 7 | TTGTACTGAAATGGCACCAG |
| Pho1a | CCCATACCAGGGTATAAGACCAG | Exon 5 | CTGGTCTTATACCCTGGTAT |

**Table S2: Genotyping primer sequences.**

| | Forward primer (5' to 3') | Reverse primer (5' to 3') |
|---|---|---|
| Pho1a Sense Fragment | CACAAGAGGACGCAACACAACACAC | CCTTAGGGAGCTCAAACCTTTCAGG |
| Pho1a Anti-sense Fragment | CACAAGAGGACGCAACACAACA | TCAAGGTATTGTTGACATGGAAACAGCG |
| SS4 Amplicon | ATTCTAGCGGCATTTCTT | ACTTGTCTTTCCACCCTT |
| SS5 Amplicon | TCTTCTGGTATATGGTTTGAATT | AAAAATGAGAGTGGCAAAAA |

**Table S3: SS4 genotyping results.** Red represents deleted bases, and blue represents inserted bases.

| Line name | Edit | Sequence |
|---|---|---|
| Wild type | | CCATATTGCAGCCGAGA |
| *ss4*-1 | 1 bp insertion | CCATATTTGCAGCCGAGA |
| | 1 bp deletion | CCATATTGCAGCCGAGA |
| | 2 bp deletion | CCATATTGCAGCCGAGA |
| *ss4*-2 | 1 bp insertion | CCATATATGCAGCCGAGA |
| | 4 bp deletion | CCATATTGCAGCCGAGA |
| | 4 bp deletion | CCATATTGCAGCCGAGA |
| | 1 bp deletion | CCATATTGCAGCCGAGA |
| *ss4*-3 | 1 bp insertion | CCATATTTGCAGCCGAGA |
| | 2 bp deletion | CCATATTGCAGCCGAGA |
| *ss4*-4 | 2 bp deletion | CCATATTGCAGCCGAGA |
| | 5 bp deletion | CCATATTGCAGCCGAGA |
| | 5 bp deletion | CCATATTGCAGCCGAGA |
| | 4 bp deletion | CCATATTGCAGCCGAGA |
| *ss4*-5 | 4 bp deletion | CCATATTGCAGCCGAGA |
| | 2 bp deletion | CCATATTGCAGCCGAGA |
| | 4 bp deletion | CCATATTGCAGCCGAGA |

**Table S4: SS5 genotyping results.** Red represents deleted bases, and blue represents inserted bases.

| **Mutant name** | **Edit** | **Sequence** |
|---|---|---|
| Wild type | | TACTGAAATGGCACCAGTGGTATCAGTTGGA |
| *ss5-1* | 2 bp deletion | TACTGAAATGGCACCAGTGGTATCAGTTGGA |
| | 5 bp deletion | TACTGAAATGGCACCAGTGGTATCAGTTGGA |
| | 8 bp deletion | TACTGAAATGGCACCAGTGGTATCAGTTGGA |
| *ss5-2* | 1 bp deletion | TACTGAAATGGCACCAGTGGTATCAGTTGGA |
| | 1 bp insertion | TACTGAAATGGCACTCAGTGGTATCAGTTGGA |
| *ss5-3* | 1 bp deletion | TACTGAAATGGCACCAGTGGTATCAGTTGGA |
| | 1 bp deletion | TACTGAAATGGCACCAGTGGTATCAGTTGGA |
| | 8 bp deletion | TACTGAAATGGCACCAGTGGTATCAGTTGGA |
| *ss5-4* | 1 bp deletion | TACTGAAATGGCACCAGTGGTATCAGTTGGA |
| | 2 bp deletion | TACTGAAATGGCACCAGTGGTATCAGTTGGA |
| | 1 bp deletion | TACTGAAATGGCACCAGTGGTATCAGTTGGA |

**Table S5: List of starch metabolic genes in potato.**

| **ADP-Glucose synthesis** | |
|---|---|
| Plastidial phosphoglucoisomerase (pPGI) | Soltu.DM.04G030730 |
| Plastidial Phosphoglucomutase (pPGM) | Soltu.DM.03G016410/Soltu.DM.03G016420 |
| ATP-ADP antiporter 1 (NTT1) | Soltu.DM.03G034530 |
| ATP-ADP antiporter 2 (NTT2) | Soltu.DM.12G010800 |
| ADP-glucose pyrophosphorylase large subunit 1 (AGPL1) | Soltu.DM.01G024440 |
| ADP-glucose pyrophosphorylase large subunit 2 (AGPL2) | Soltu.DM.07G010140 |
| ADP-glucose pyrophosphorylase large subunit 3 (AGPL3) | Soltu.DM.01G049590 |
| ADP-glucose pyrophosphorylase small subunit 1.1 (AGPS1.1) | Soltu.DM.07G022290 |
| ADP-glucose pyrophosphorylase small subunit 1.2 (AGPS1.2) | Soltu.DM.12G028820 |
| ADP-glucose pyrophosphorylase small subunit 2 (AGPS2) | Soltu.DM.08G006240 |
| **Granule Initiation proteins** | |
| Protein Targeting to Starch 2a (PTST2a) | Soltu.DM.05G006480 |
| Protein Targeting to Starch 2b (PTST2b) | Soltu DM.01G008470 |
| Protein Targeting to Starch 3 (PTST3) | Soltu.DM.09G005340 |
| MAR-Binding Filament Protein (MFP1) | Soltu.DM.03G034880 |
| Myosin-Resembling Chloroplast Protein (MRC) | Soltu.DM.08G009720 |
| **Amylopectin Synthesis** | |
| Starch Synthase I (SS1) | Soltu.DM.03G022350 |

| | |
|---|---|
| Starch Synthase II (SS2) | Soltu.DM.02G031690 |
| Starch Synthase III (SS3) | Soltu.DM.02G020170 |
| Starch Synthase IV (SS4) | Soltu.DM.02G014060 |
| Starch Synthase V (SS5) | Soltu.DM.02G027020 |
| Starch Synthase VI (SS6) | Soltu.DM.07G013630 |
| Starch branching enzyme 1 (SBE1) | Soltu.DM.04G037620 |
| Starch branching enzyme 2 (SBE2) | Soltu.DM.09G004100 |
| Isoamylase 1.1 (ISA1) | Soltu.DM.07G005540 |
| Isoamylase 1.2 (ISA 1.2) | Soltu.DM.10G007060 |
| Isoamylase 2 (ISA2) | Soltu.DM.09G019230 |
| Early Starvation 1a (ESV1a) | Soltu.DM.04G037500 |
| Early Starvation 1b (ESV1b) | Soltu.DM.12G004870 |
| Like Early Starvation (LESV) | Soltu.DM.06G014960 |
| **Amylose Synthesis** | |
| Granule bound starch synthase 1 (GBSS1) | Soltu.DM.08G030230 |
| Protein Targeting to Starch 1 (PTST1) | Soltu.DM.02G026830 |
| **MOS Metabolism** | |
| Alpha-glucan phosphorylase 1a-1 (PHO1a-1) | Soltu.DM.03G007710/Soltu.DM.03G007720 |
| Alpha-glucan phosphorylase 1a-2 (PHO1a-2) | Soltu.DM.03G007750/Soltu.DM.03G007760 |
| Alpha-glucan phosphorylase 1b (PHO1b) | Soltu.DM.05G000570 |
| Alpha-glucan phosphorylase 2 (PHO2) | Soltu.DM.09G011580 |
| Alpha-glucan phosphorylase 3 (PHO3) | Soltu.DM.02G017070 |
| Disproportionating enzyme 1 (DPE1) | Soltu.DM.04G022990 |
| Disproportionating enzyme 2 (DPE2) | Soltu.DM.02G000530 |
| **Reversible Starch Phosphorylation** | |
| Glucan water dikinase (GWD) | Soltu.DM.05G009520 |
| Phosphoglucan water dikinase (PWD) | Soltu.DM.09G030970 |
| Phosphoglucan phosphatase (SEX4a) | Soltu.DM.11G004900 |
| Phosphoglucan phosphatase (SEX4b) | Soltu.DM.03G024560 |
| Like SEX4 1 (LSF1) | Soltu.DM.12G016610 |
| Like SEX4 2 (LSF2) | Soltu.DM.06G010900 |
| **Starch Degradation** | |
| Alpha-amylase 1.1 (AMY1) | Soltu.DM.04G033700 |
| Alpha-amylase 1.2 (AMY1.2) | Soltu.DM.03G013410 |
| Alpha-amylase 2 (AMY23) | Soltu.DM.04G037250 |
| Alpha-amylase 3 (AMY3) | Soltu.DM.05G006330 |
| Alpha-amylase 4 (AMY4) | Soltu.DM.02G009320 |
| Beta-amylase 1 (BAM1) | Soltu.DM.09G027770 |
| Beta-amylase 3.1 (BAM3.1) | Soltu.DM.08G001120 |
| Beta-amylase 3.2 (BAM3.2) | Soltu.DM.08G023420 |

| | |
|---|---|
| Beta-amylase 5 (BAM5) | Soltu.DM.07G018100 |
| Beta-amylase 7 (BAM7) | Soltu.DM.01G033560 |
| Beta-amylase 8 (BAM8) | Soltu.DM.08G003130 |
| Beta-amylase 9 (BAM9) | Soltu.DM.01G022570 |
| Beta-amylase 10 (BAM10) | Soltu.DM.08G029750 |
| Isoamylase 3 (ISA3) | Soltu.DM.06G000410 |
| Limit dextrinase (LDA) | Soltu.DM.11G004600 |
| **Plastidial sugar transporters** | |
| Glucose transporter (GLT1) | Soltu.DM.02G026040 |
| Glucose-6-phosphate translocator 1.1 (GPT1) | Soltu.DM.07G025810 |
| Glucose-6-phosphate translocator 2.2 (GPT2) | Soltu.DM.07G017870 |
| Glucose-6-phosphate translocator 2.1 (GPT2.1) | Soltu.DM.05G018750 |
| Maltose excess 1 (MEX1) | Soltu.DM.04G025730 |
| Triose-phosphate/phosphate translocator (TPT) | Soltu.DM.10G004860 |
| Triose-phosphate/phosphate translocator-like (TPT-like) | Soltu.DM.01G008290 |
| **Sucrose metabolism** | |
| Sucrose synthase (SUSIa) | Soltu.DM.07G013360 |
| Sucrose synthase (SUSIb) | Soltu.DM.07G013370 |
| Sucrose synthase (SUSIc) | Soltu.DM.12G026390 |
| Sucrose synthase (SUSII) | Soltu.DM.09G031820 |
| Sucrose synthase (SUSIIIa) | Soltu.DM.02G020800 |
| Sucrose synthase (SUSIIIb) | Soltu.DM.03G019120 |
| UDP-glucose pyrophosphorylase 1 (UGPase1) | Soltu.DM.11G001030 |
| UDP-glucose pyrophosphorylase 2 (UGPase2) | Soltu.DM.05G024440 |
| UDP-glucose pyrophosphorylase 3 (UGPase3) | Soltu.DM.01G028790 |
| Sucrose Phosphate Synthase 1 (SPSI) | Soltu.DM.07G003160 |
| Sucrose Phosphate Synthase 2 (SPSII) | Soltu.DM.08G010240 |
| Sucrose Phosphate Synthase 3 (SPSIII) | Soltu.DM.09G029230 |
| Sucrose Phosphate Synthase 4 (SPSIV) | Soltu.DM.11G017190 |
| Hexokinase 1 (HXK1) | Soltu.DM.07G001270 |
| Hexokinase 2 (HXK2) | Soltu.DM.03G035820 |
| Hexokinase 3 (HXK3) | Soltu.DM.12G025470 |
| Hexokinase 4 (HXK4) | Soltu.DM.04G036520 |
| Hexokinase 5 (HXK5) | Soltu.DM.11G019290 |
| Hexokinase 6 (HXK6) | Soltu.DM.02G027090 |
| Fructokinase 1 (FRK1) | Soltu.DM.03G016430 |

| | |
|---|---|
| Fructokinase 2 (FRK2) | Soltu.DM.06G028540 |
| Fructokinase 3 (FRK3) | Soltu.DM.02G026790 |
| Fructokinase 4 (FRK4) | Soltu.DM.10G006860 |
| Fructokinase 5 (FRK5) | Soltu.DM.11G016260 |
| Cytosolic phosphoglucomutase (cPGM) | Soltu.DM.04G015010 |
| Cytosolic phosphoglucoisomerase (cPGI) | Soltu.DM.12G030200 |
| Vacuolar Invertase (vINV) | Soltu.DM.03G015280 |
| Cell wall Invertase 1 (cwINV1) | Soltu.DM.03G036480 |
| Cell wall Invertase 2 (cwINV2) | Soltu.DM.08G008790 |
| Cell wall Invertase 3 (cwINV3) | Soltu.DM.06G021110 |
| Invertase 1 (INV1) | Soltu.DM.08G025360 |
| Invertase 2 (INV2) | Soltu.DM.09G002900 |
| Invertase 3 (INV3) | Soltu.DM.10G025030 |
| Invertase 4 (INV4) | Soltu.DM.10G022430 |
| Invertase 5 (INV5) | Soltu.DM.10G021970 |
| Invertase 6 (INV6) | Soltu.DM.09G002890 |
| Invertase 7 (INV7) | Soltu.DM.10G025040 |
| Neutral Invertase 1 (NINV1) | Soltu.DM.01G018690 |
| Neutral Invertase 2 (NINV2) | Soltu.DM.01G040550 |
| Neutral Invertase 3 (NINV3) | Soltu.DM.11G021450 |
| Neutral Invertase 4 (NINV4) | Soltu.DM.11G006090 |
| Neutral Invertase 5 (NINV5) | Soltu.DM.06G020260 |
| Neutral Invertase 6 (NINV6) | Soltu.DM.11G012260 |
| Neutral Invertase 7 (NINV7) | Soltu.DM.04G036560 |
| Neutral Invertase 8 (NINV8) | Soltu.DM.01G050680 |
| **Other** | |
| Branching enzyme 3a (SBE3a/MKIP1a) | Soltu.DM.07G026510 |
| Branching enzyme 3b (SBE3b/MKIP1b) | Soltu.DM.07G025710 |
| Putative Phosphoglucomutase (PGM) | Soltu.DM.07G015450/Soltu.DM.07G015460 |
